# Host specificity, seasonality, and urbanicity shape the gut microbiome of a synanthropic insect

**DOI:** 10.64898/2026.10.28.762591

**Authors:** Hanna Noyes, Nur-E-Jannat Anika, Julia Combs, Tamheeda Khan, Josefine Just, Su Fitzpatrick, Rowan Graham Angelini, David R. Angelini

## Abstract

The soapberry bug *Jadera haematoloma* is a classic model of rapid adaptive evolution, with a well-documented host shift from native herbaceous sapinds to the introduced goldenrain tree (*Koelreuteria spp.*). This host shift has facilitated the insect’s spread into novel anthropogenic urban habitats. Microbial associations are often critical to insect environmental adaptation. Here, we characterize the gut microbiota of *J. haematoloma* across native and introduced host plants, seasonal transitions, and a gradient of urbanicity. Our findings indicate that the soapberry bug gut microbiome is highly specialized, with a small number of core bacterial amplicon sequence variants (ASVs) primarily from the families Rhizobiaceae, Streptococcaceae, Enterobacteriaceae, and Lactobacillaceae. While traits such as sex and wing morph do not correlate with microbiome structure, the host plant acts as the primary determinant of community composition. Bugs feeding on the ancestral host plant maintain a more diverse and even microbial community featuring *Enterobacteriaceae*, whereas the gut environment of bugs on the introduced host is overwhelmingly dominated by a single unclassified *Rhizobiaceae* ASV. Furthermore, seasonal transitions drive significant turnover among accessory microbial taxa. Finally, we demonstrate that urbanization introduces a distinct ecological effect on the insect’s microbiome. While the degree of local green space does not systematically alter within-host taxonomic richness, it is associated with modest change in mean community composition and significantly disrupts microbial community variance. Increased urbanization correlates with significantly higher beta dispersion, indicating that anthropogenic environments lead to more stochastic microbiomes, which may be an indicator of environmental stress. Together, these results illustrate how rapid dietary host shifts and the ecological pressures of urbanization interact to shape the microbiome of a synanthropic insect.

## Introduction

The soapberry bug *Jadera haematoloma* (Hemiptera: Rhopalidae) is a classic model of rapid ecological adaptation to synanthropic (human-dominated) environments. Native to coastal hammock forests from the Gulf of Mexico to South America, these insects are specialized to feed on plants of the soapberry family, which are rich in saponins and cyanolipids. In southern Florida, the most common historical host plants were perennial *Cardiospermum* spp. vines. However, since the mid-20th century, populations of *J. haematoloma* have undergone a rapid host shift, adapting to the introduced goldenrain tree (*Koelreuteria* spp.). This transition presents the bugs with distinctly different nutritional and phenological environments; while *Cardiospermum* provides relatively low but consistent seed production, *Koelreuteria* trees produce numerous, lipid-rich seeds during a restricted seasonal window. This host shift has driven well-documented, rapid divergence in insect morphology, life history, and nutritionally regulated flight-fecundity polyphenism (Carroll and Boyd 1992; Fawcett et al. 2018).

Utilization of the introduced host plant of *J. haematoloma* has facilitated the insect’s spread into novel anthropogenic habitats. Because goldenrain trees were widely planted as ornamentals in urban and suburban landscapes during the mid-to-late twentieth century, derived ecotypes of *J. haematoloma* have been exposed to the unique ecological pressures of urbanization, such as altered microclimates, habitat fragmentation, and pollution.

The gut microbiome is increasingly recognized as a critical ecological factor, often expanding the metabolic capabilities of the insect, facilitating dietary shifts, and buffering against environmental stress (Engel & Moran 2013; Douglas 2015; Young et al. 2025). In many hemipterans, microbial symbionts synthesize essential nutrients missing from restricted diets (Gündüz & Douglas 2009) or assist in detoxifying plant defensive compounds (Itoh et al 2018). Furthermore, environmental stressors such as extreme seasonal fluctuations and urbanization have been shown to significantly alter host-associated microbiomes in various taxa, sometimes destabilizing the community structure (Ferguson et al. 2018; Maraci et al. 2022). Given the rapid evolutionary divergence and ecological expansion of *J. haematoloma*, characterizing its microbial associations is an important step in understanding its adaptation to the novel anthropogenic environment.

Here, we characterize the gut microbiota of *J. haematoloma* across native and introduced host plants, seasonal transitions, and a local gradient of urbanicity. We show that the global gut community is heavily structured by a small number of core bacterial amplicon sequence variants (ASVs). While sex and wing morph do not correlate with the microbiome, we find that host plant is the primary determinant of community composition, alongside significant seasonal turnover of accessory taxa. Finally, we demonstrate that while urbanicity does not systematically shift within-host diversity, it significantly disrupts the microbial community through increased beta dispersion, which is characteristic of environmental stress (Zaneveld et al. 2017).

## Results

### The soapberry bug microbiome is dominated by a small number of core taxa

Soapberry bugs were collected from 13 sites in southern Florida (Figure S1; Table S1) in January and July 2019. Most specimens were found on goldenrain trees (*Koelreuteria paniculata*), however July samples included individuals from the native host plant, *Cardiospermum corindum*. To characterize the gut microbiome of *J. haematoloma*, we sequenced the 16S V6-V8 region. Following quality filtering and read merging, sequence variants were denoised and tabulated (Table S2). After the removal of chloroplasts, mitochondria, and filtering for minimum library size thresholds, our final dataset comprised 144 individual samples containing a total of 2,064 distinct ASVs.

Notably, initial *in silico* taxonomic classification identified a substantial presence of the widespread intracellular bacterium *Wolbachia*. Since this endosymbiont would skew compositional data intended to describe the gut microbiome, all *Wolbachia*-assigned ASVs were filtered out of the dataset for all downstream diversity and differential abundance analyses.

To confirm that *Wolbachia* reads are consistently associated with the tissues of *J. haematoloma* we independently screened somatic tissues. The head, gut and gonads from four females and four males, as well as three pools of eggs, were used as starting templates in PCR reactions with two *Wolbachia*-specific primer pairs. All insect-derived samples produced products in one or both reactions, while water controls remained negative (Figure S1). This suggests that, as in other insects, *Wolbachia* is an intracellular endosymbiont of *J. haematoloma*.

The *J. haematoloma* gut microbiota composition overall was dominated by two bacterial phyla: Proteobacteria and Firmicutes. At the family level, the global community was heavily structured by just four dominant lineages, which together accounted for nearly 96% of the total mean relative abundance: Rhizobiaceae (67.02%), Streptococcaceae (14.78%), Enterobacteriaceae (9.55%), and Lactobacillaceae (4.54%).

We defined the core microbiome for each host plant-associated cohort, applying thresholds of 80% prevalence and 10% mean relative abundance (Table S3). In the *Cardiospermum corindum* cohort, the core microbiome consisted of four ASVs, which were present in all of the sampled individuals: two unclassified Enterobacteriaceae ASVs (mean abundances of 27.9% and 23.0%), an unclassified Alphaproteobacteria (24.2%), and a *Lactococcus* ASV (15.0%). In contrast, the *Koelreuteria paniculata* core community was restricted to just two taxa, dominated by a single unclassified Rhizobiaceae that accounted for 66.2% of the mean relative abundance and was present in 99.3% of the samples. Notably, the second core member of the *K. paniculata* cohort was the same *Lactococcus* sequence variant (mean abundance 13.5%, prevalence 84.2%) identified in the *C. corindum* core, making it the only globally conserved core ASV across both host plant populations. Three of the other top-five most abundant ASVs in *C. corindum* are present at much lower mean relative abundances in *K. paniculata* samples (Figure 2B).

**Figure 1.**
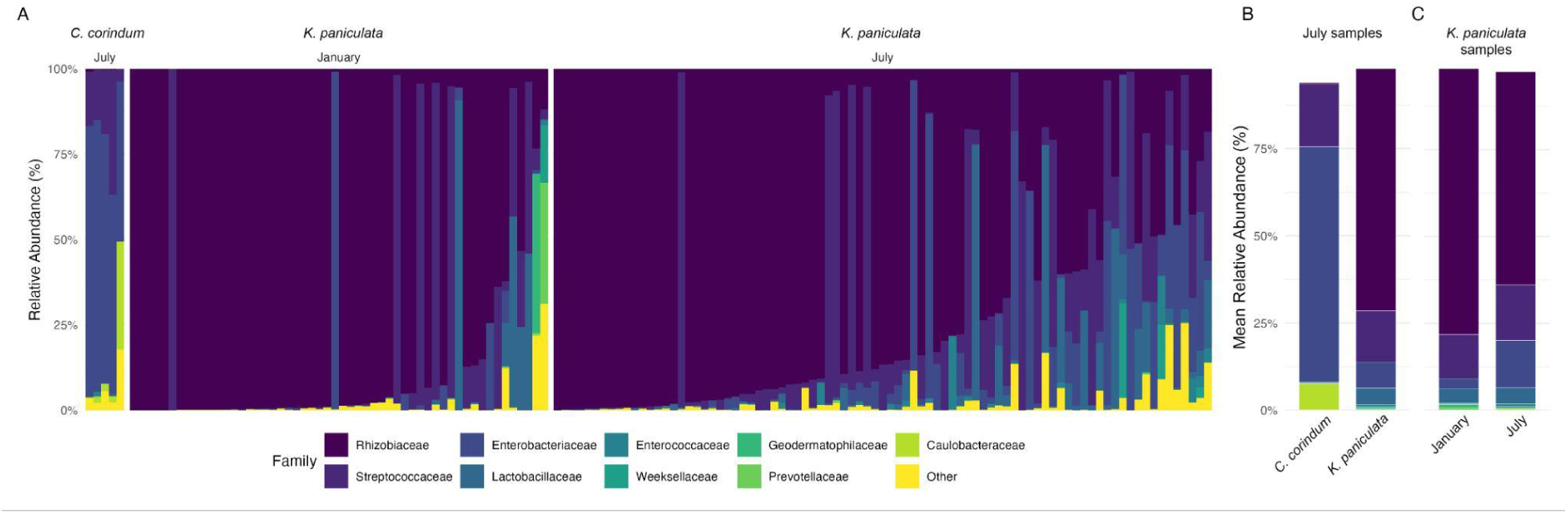
Taxonomic composition of the *Jadera haematoloma* gut microbiota. Relative abundance of bacterial taxa at the family level based on 16S rRNA gene amplicon sequencing, following the exclusion of the endosymbiont *Wolbachia*. Stacked barplots illustrate the global family-level composition based on 144 individual samples, highlighting the dominance of Rhizobiaceae, Streptococcaceae, Enterobacteriaceae, and Lactobacillaceae. Taxa with a mean relative abundance of less than 1% across the dataset are grouped as “Other.” **(A)** Individual samples, grouped by host plant and month of collection. **(B)** Mean relative abundance of bacterial families partitioned by host plant, comparing the gut microbiota of bugs feeding on balloon vine (*Cardiospermum corindum*; BV) versus goldenrain tree (*Koelreuteria paniculata*; GRT). **(C)** Mean relative abundance of bacterial families partitioned by season, comparing January and July sample cohorts.

**Figure 2.**
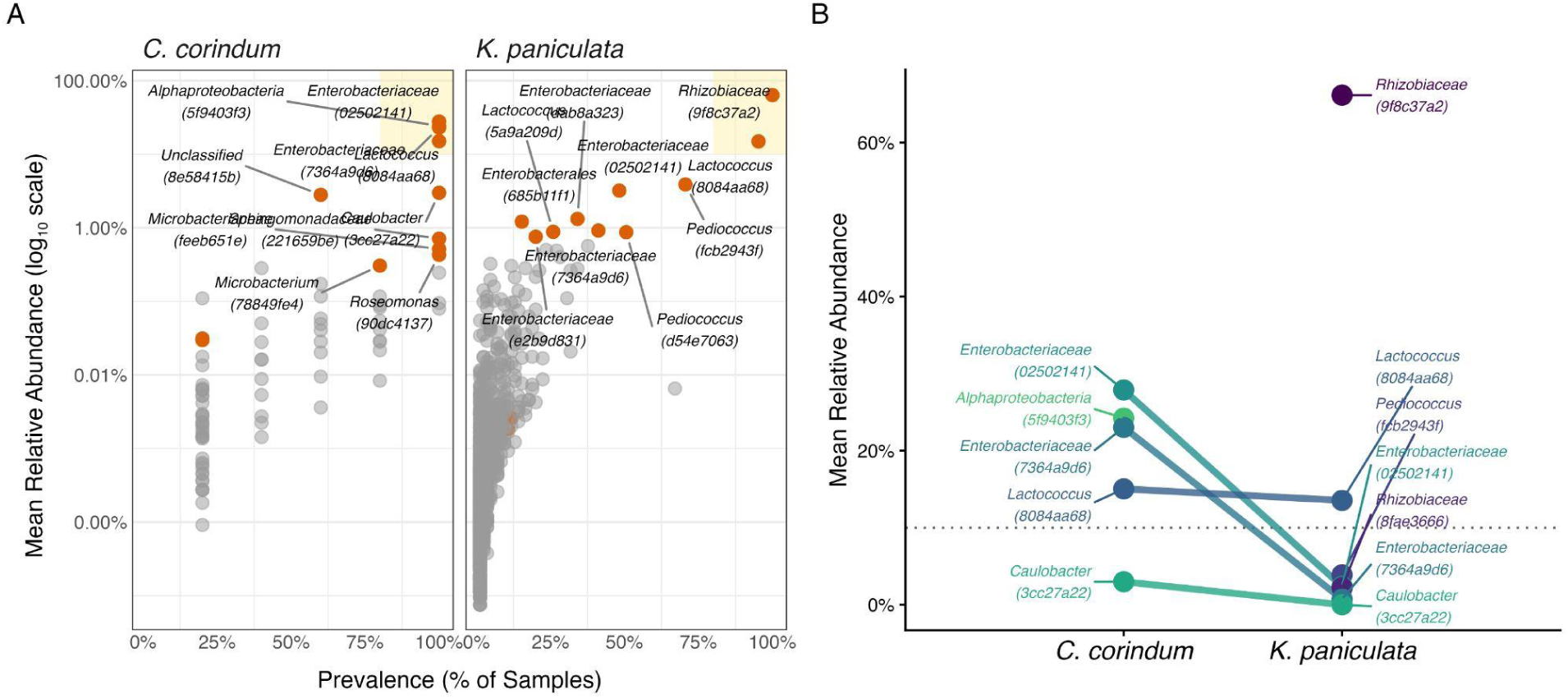
Core microbiome differences by host plant. **(A)** Scatterplot of the prevalence of ASVs in *C. corindum* (left) and *K. paniculata* (right) host plants against log_10_ mean relative abundance. The yellow box highlights ASVs with high prevalence and mean relative abundance defined as core microbiota. The ten most abundant ASVs in each host plant cohort are named and highlighted. **(B)** Slope plot comparing mean relative abundance of the five most abundant ASVs between *C. corindum* and *K. paniculata*. The dotted line indicates the 10% mean relative abundance threshold for core microbiota. Note that the unclassified Alphaproteobacteria ASV in *C. corindum* samples and the unclassified Rhizobiaceae ASV in *K. paniculata* are endemic to those cohorts.

Interestly, each host plant cohort is dominated by endemic ASVs. The unclassified Alphaproteobacteria in *C. corindum* does not appear in *K. paniculata* samples and the unclassified Rhizobiaceae that dominates *K. paniculata* gut communities is not found in *C. corindum* samples.

### Microbial diversity within individuals is influenced by seasonality and host plant

To assess the within-sample taxonomic richness and evenness of the *J. haematoloma* gut microbiota, we calculated observed ASVs (richness), alongside the Shannon and Simpson diversity indices. Individual sample metrics and comparative statistics are available in the supplemental materials (Table S4, Table S5). We first evaluated intrinsic host traits and found that neither sex nor wing morphology significantly influenced alpha diversity across any of the three measured metrics (Figure S2A-B; Wilcoxon rank-sum tests, p > 0.05), indicating that these adult phenotypes do not strongly dictate gut microbiome structure.

In contrast, environmental factors exerted distinct effects on within-sample diversity (Figure 3). When comparing host plant within the July cohort, absolute community richness (observed ASVs) did not differ significantly between bugs feeding on *C. corindum* versus *K. paniculata* (W = 316.0, p = 0.069). However, the evenness and proportional diversity of the communities were significantly altered by the host plant, as captured by the Shannon (W = 352.0, p = 0.014) and Simpson indices (W = 356.0, p = 0.012) (Figure 3A). Seasonality was also associated with taxonomic diversity among the *K. paniculata* cohort; comparing January to July revealed highly significant differences in community richness (W = 1491.0, p = 0.0005) as well as Shannon (W = 1306.5, p < 0.0001) and Simpson diversity (W = 1337.5, p < 0.0001) (Figure 3B).

**Figure 3.**
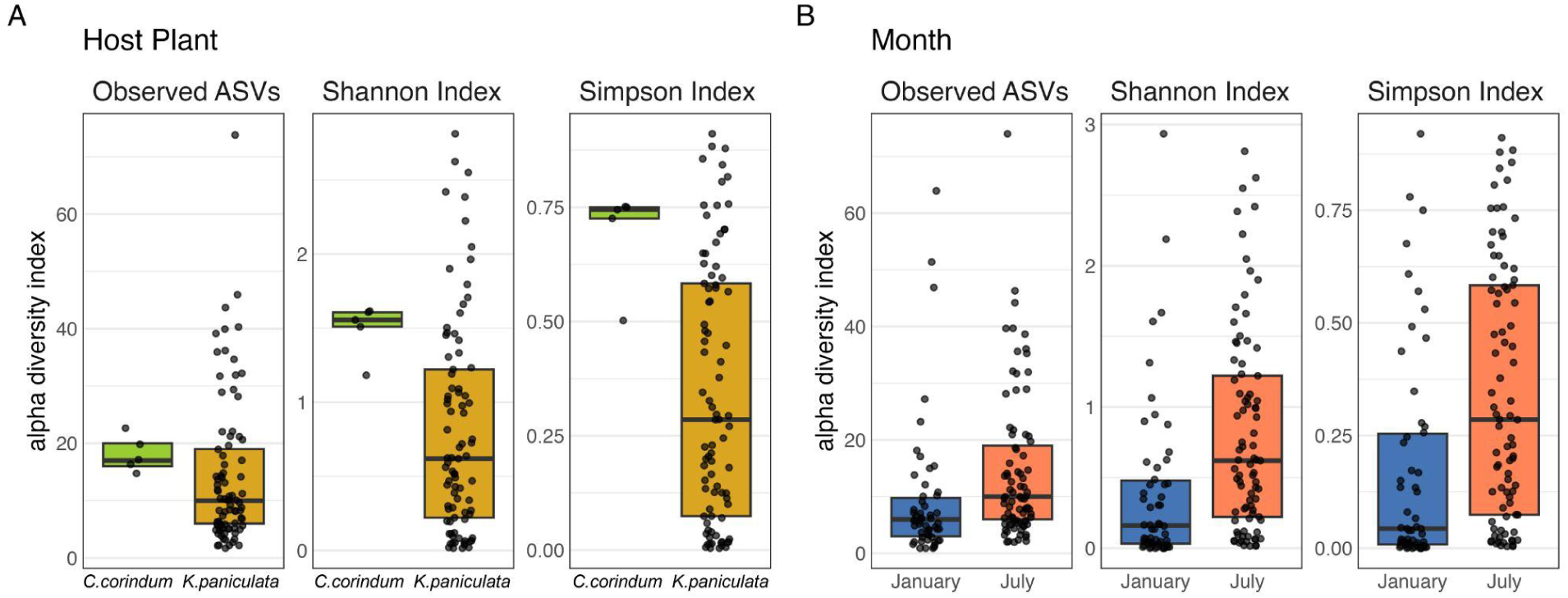
Alpha diversity of the *Jadera haematoloma* gut microbiota is significantly influenced by host plant and seasonality. Box plots displaying within-sample taxonomic richness and diversity. Each panel plots observed ASVs, Shannon index and Simpson’s index. Center lines represent the median, box limits indicate the upper and lower quartiles, and whiskers extend to 1.5 times the interquartile range. **(A)** Alpha diversity comparisons between host plant diets within the July sample cohort. Bugs feeding on *Cardiospermum corindum* (BV, n = 5) versus *Koelreuteria paniculata* (GRT, n = 85) exhibit significant differences in community evenness (Shannon and Simpson indices, p < 0.05). **(B)** Alpha diversity comparisons between seasons (January: n = 54; July, n = 85) within the *K. paniculata* cohort. All three metrics demonstrate significant seasonal shifts in microbiome richness and diversity (Table S5). Statistical significance was determined via Wilcoxon rank-sum tests. Box plots for non-significant host and landscape variables (sex, wing morph, urbanization and green space) are provided in Figure S3.

Finally, to test whether continuous landscape variables broadly alter within-sample diversity, we utilized linear mixed models incorporating the collection site as a random intercept. Unlike host plant and seasonality, neither the degree of surrounding green space nor urbanicity significantly predicted alpha diversity across any metric (p > 0.17 for all comparisons). This suggests that urbanization does not systematically expand or collapse the absolute number or evenness of taxa residing within individual insects.

### Community structure is predominantly associated with the host plant

To assess broad-scale patterns in the *J. haematoloma* gut microbiota, we first evaluated global community variance using principal coordinates analysis (PCoA) on distance matrices describing community composition (Bray-Curtis), presence/absence of phylogenetic lineages (unweighted UniFrac), and the abundance-weighted phylogenetic distance (weighted UniFrac). Separation of host plant-associated samples into distinct clusters in the Bray-Curtis PCoA suggested different dominant ASV in these groups (Figure 4C). However, with unweighted and weighted UniFrac distances, *C. corindum* samples nested within a larger space occupied by *K. paniculata* samples (Figure S4B-C,H-I). This nested relationship suggests that *K. paniculata* supports a wider range of microbial lineages. In contrast, samples did not separate by collection month, displaying broader overlap (Figure S4D-F, J-L). This overlap suggests an overall stability across seasons. We noted that July samples were more clustered than January samples in the unweighted UniFrac plot (Figure 4D), suggesting that rare lineages in January may be outcompeted by a smaller number of lineages in July.

**Figure 4.**
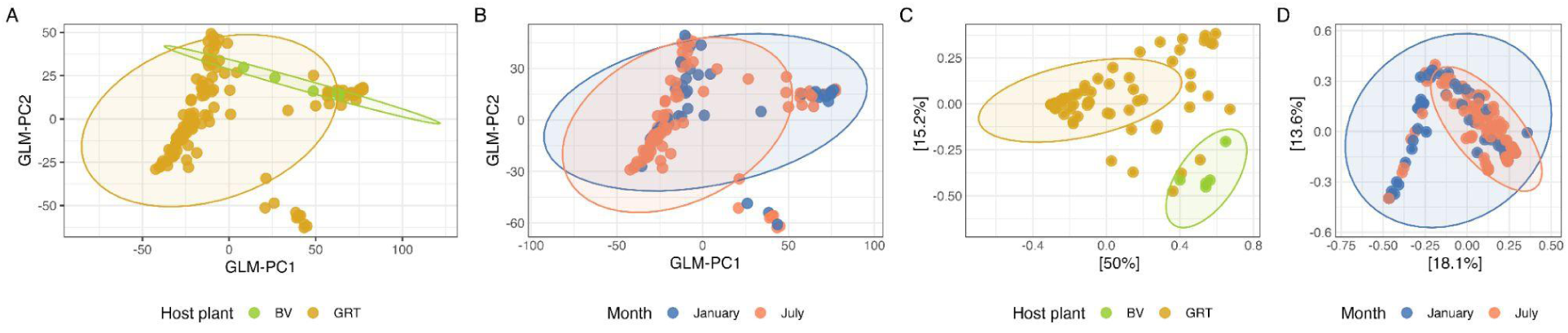
Global community structure and drivers of the *Jadera haematoloma* gut microbiota. **(A–B)** Generalized linear model principal component analysis (GLM-PCA) ordination plots of the global dataset (n = 144) under a negative binomial error distribution, illustrating community clustering. **(A)** GLM-PCA colored by host plant species (*C. corindum*, BV vs. *K. paniculata*, GRT) and **(B)** sampling month (January vs. July). **(C)** Principal coordinates analysis (PCoA) based on Bray-Curtis dissimilarity within the July sample cohort (n = 90), demonstrating significant community separation by host plant. **(D)** PCoA based on unweighted UniFrac distance within the *K. paniculata* (GRT) cohort (n = 139). The nested distribution of July samples within the broader January variance highlights a seasonal restriction in phylogenetic lineage diversity rather than complete community turnover. Shaded ellipses represent 95% confidence intervals assuming a multivariate distribution (stat_ellipse). Full PERMANOVA model outputs corresponding to panels C and D are detailed in Table S6.

We also used generalized linear model principal component analysis (GLM-PCA) to examine community structure. This approach accounts for overdispersion in the count data that often characterizes microbiome datasets. GLM-PCA also demonstrated a nesting of the microbiota associated with the ancestral host plant *C. corindum* within the larger community-space defined by *K. paniculata* microbiomes. (Figure 4A) and a more limited separation based on the month of collection (Figure 4B).

Initial global permutational multivariate analysis of variance (PERMANOVA) models supported these trends, identifying both host plant and season as significant factors in community variance (Table S6). However, because field sampling was unbalanced, with *C. corindum* sampled only in July, these global patterns may be confounded. Therefore, to better identify ecological signals, subsequent PERMANOVA was restricted to data subsets. We evaluated the influence of host plant within the July sample cohort, and the influence of seasonal shifts within the *K. paniculata* (goldenrain tree) cohort. While PERMANOVA is fairly robust to differences in sample sizes, unequal group variances can increase the false positive error rate. We tested for differences in multivariate beta dispersion among distance matrices for all categorical factors. Only unweighted UniFrac showed a significant difference by month (betadisper, p = 0.002), which was greater in January samples compared to July.

Within the July cohort, host plant species had a significant influence on gut community structure. This dietary effect was robust across three beta diversity metrics, driving significant shifts in community composition (Bray-Curtis: R^2^ = 0.14451, p < 0.001) (Figure 4C), presence/absence of phylogenetic lineages (unweighted UniFrac: R^2^ = 0.07362, p < 0.001), and the abundance-weighted phylogenetic core (weighted UniFrac: R^2^ = 0.03447, p = 0.04595) (Table S6). Holding the host plant constant within the *K. paniculata* cohort, the transition from January to July appears to somewhat stabilize the accessory microbiome. While seasonal shifts did not significantly alter the abundance-weighted taxonomic structure (Bray-Curtis: R^2^ = 0.015, p = 0.104) or the dominant microbial lineages (weighted UniFrac: R^2^ = 0.005, p = 0.424), they drove a highly significant shift in unweighted UniFrac distances (R^2^ = 0.041, p < 0.001) (Table S6). This analysis helps explain the nesting of July samples concentrated within one area of the broader variance of the January samples (Figure 4D).

### Consensus modeling identifies taxonomic drivers of host plant and seasonal community differences

To identify the specific bacterial lineages associated with observed differences in the beta diversity models, we applied a multi-algorithm consensus approach using ANCOM-BC2 (Table S7), ALDEx2 (Table S8), and MaAsLin3 (Table S9). To control for the methodological biases inherent to different compositional algorithms, taxa were primarily considered robust biological indicators if they achieved statistical significance (adjusted p-value < 0.05) in at least two of the three methods.

Within the July cohort, this consensus approach identified one ASV that was significantly differentially abundant between host plant populations (Figure 5A). This *Caulobacter* ASV was significantly enriched in the gut microbiota of bugs feeding on *C. corindum* relative to *K. paniculata* (Table S10; ALDEx2 effect size = −2.67, adjusted p = 0.038). When evaluating the effect of seasonality within the *K. paniculata* cohort, the consensus models identified two ASVs that differed significantly between January and July. The gut community in July was characterized by the significant enrichment of two species of Bacilli (Table S10), *Pediococcus* (ALDEx2 effect size = 0.74, FDR < 0.001) and *Lactococcus* (ALDEx2 effect size = 0.599, adjusted p = 0.0045). This *Lactococcus* ASV is also a member of the core microbiome (Table S3).

**Figure 5.**
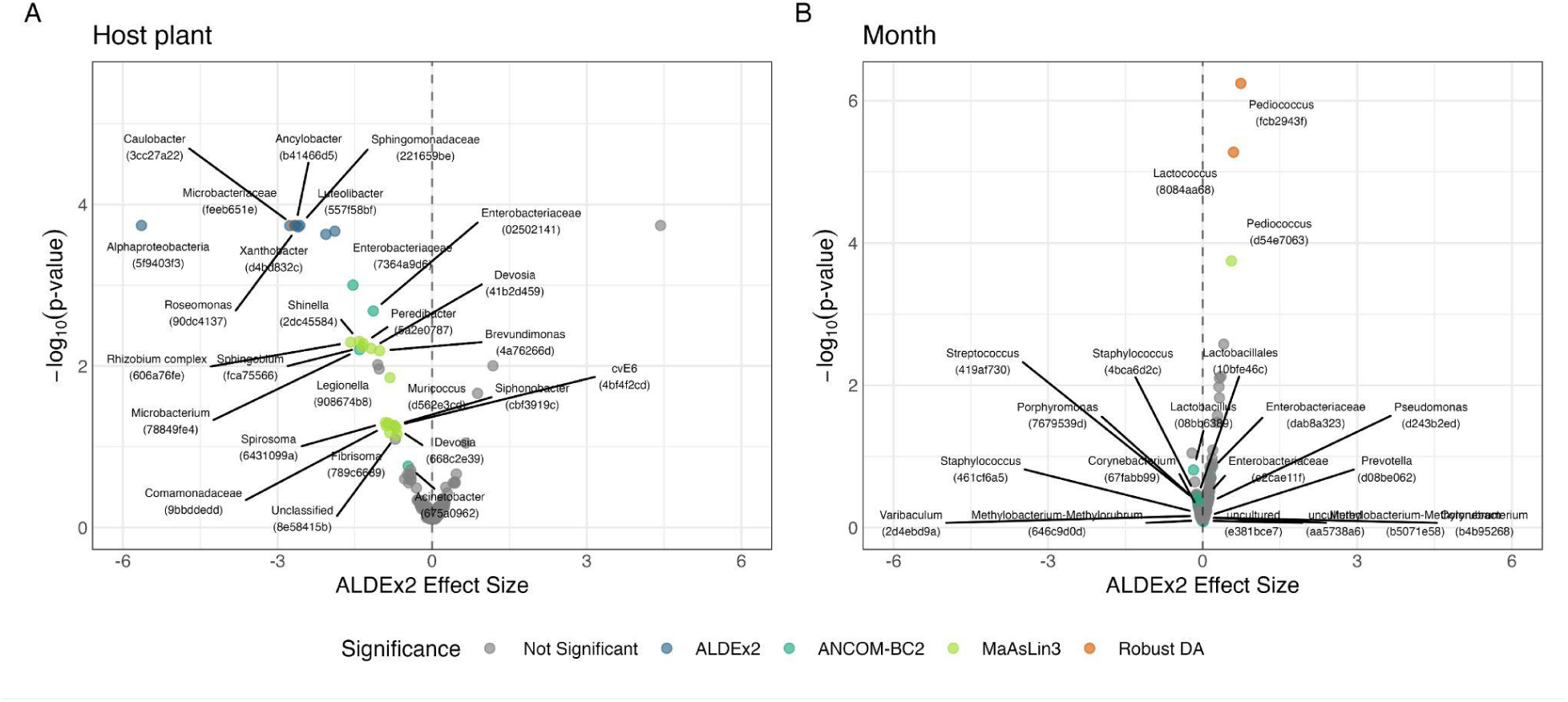
Consensus differential abundance of *Jadera haematoloma* gut microbiota driven by host plant and seasonality. Volcano plots displaying the magnitude and significance of differentially abundant ASVs as modeled by ALDEx2. The horizontal axis represents the ALDEx2 effect size, and the vertical axis represents the -log_10_ p-value. Grey points represent background ASVs, while colored, labeled points highlight taxa identified via each differential abundance analysis method (ANCOM-BC2, ALDEx2, and MaAsLin3). Robust DA taxa are defined here as those identified by at least two DA analysis methods **(A)** Differential abundance between host plants within the July sample cohort. Negative estimates indicate significant taxonomic enrichment in bugs collected on *C. corindum* relative to those from *K. paniculata*. **(B)** Differential abundance between seasons within the *K. paniculata* sample cohort. Negative estimates indicate significant taxonomic enrichment in January relative to July samples.

**Figure 6.**
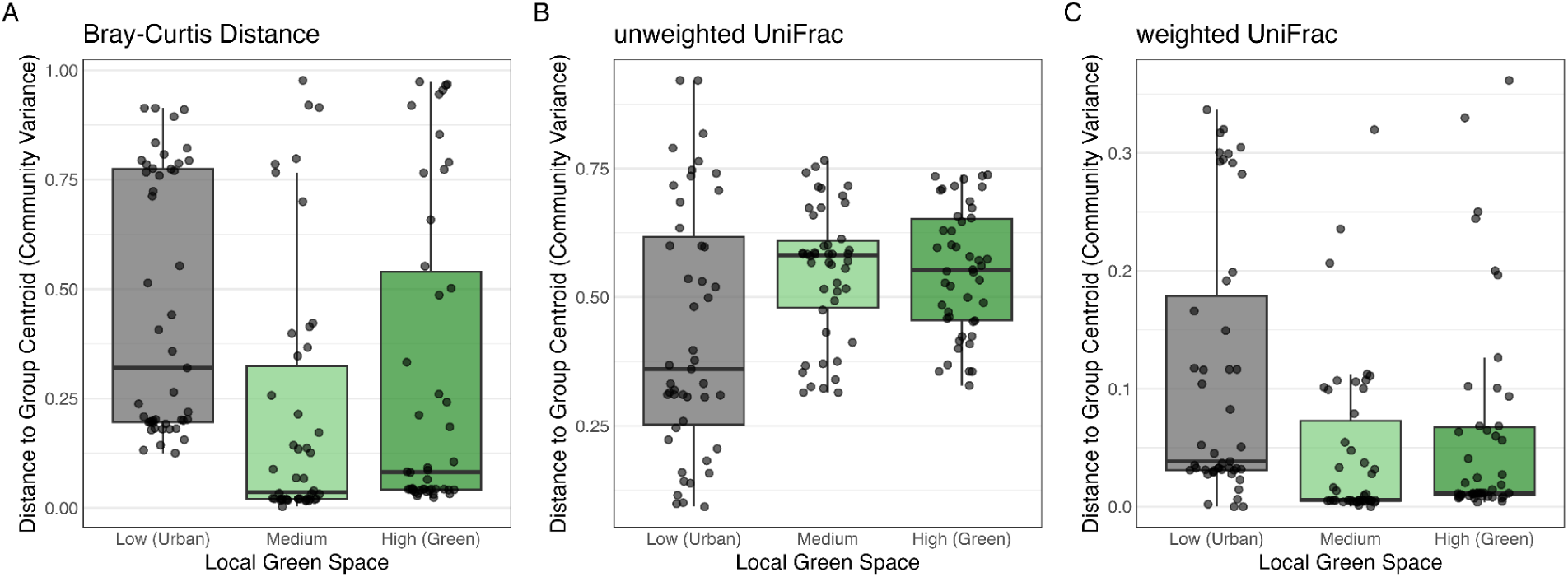
Urbanization correlates with altered beta dispersion in the *J. haematoloma* gut microbiota. Bar plots displaying the multivariate dispersion of samples, measured as the distance of individual samples to their group centroid, across an urbanization gradient. Samples are binned into low, medium, and high tertiles based on the degree of surrounding green space. Panels represent dispersion calculated using **(A)** Bray-Curtis dissimilarity, **(B)** unweighted UniFrac, and **(C)** weighted UniFrac distance metrics. Significant differences in multivariate dispersion (PERMDISP) were observed across all three metrics (Table S11).

### Urbanization correlates with compositional shifts and altered beta dispersion

To investigate the potential effects of urbanization on the *J. haematoloma* gut microbiota, we analyzed community structure across a gradient of local green space (Nguyen & Rehan 2022; Borden et al. 2025). Treating this environmental gradient as a continuous variable, global PERMANOVA models indicated that the degree of green space exerted a statistically significant, albeit modest, effect on overall community composition (Bray-Curtis: R^2^ = 0.02937, p = 0.001998; unweighted UniFrac: R^2^ = 0.0226, p = 0.000999; weighted UniFrac: R^2^ = 0.025, p = 0.04296).

Beyond shifts in average community composition, we tested whether urbanization influenced the stability and variability of the microbiome using tests for homogeneity of multivariate dispersions (PERMDISP). This analysis revealed that the degree of surrounding green space was significantly associated with altered community variance. This dispersion effect was highly robust, driving significant differences in multivariate spread across abundance-weighted taxonomic metrics (Bray-Curtis: F = 7.33, p = 0.001), as well as both presence-absence (unweighted UniFrac: F = 7.09, p = 0.002) and abundance-weighted (weighted UniFrac: F = 6.37, p = 0.002) phylogenetic distances (Table S11). Unlike the pronounced visual clustering observed for host plants, the influence of urbanization manifested as a subtle effect on both the community centroid and multivariate variance, acting on the microbiota without driving the samples into visually isolated ordination clusters.

## Discussion

### The soapberry bug gut core microbiome

The gut microbiome of *Jadera haematoloma* is dominated by a small group of bacterial taxa. Across all samples, the community is dominated by just four bacterial families, Rhizobiaceae, Streptococcaceae, Enterobacteriaceae, and Lactobacillaceae, which together account for 96% of the filtered sequencing reads (Figure 1). This extreme dominance suggests a highly specialized environment and microbial community within the soapberry bug gut. Similarly limited microbial communities have been found in other Hemiptera, where a small number of symbionts facilitate specific metabolic functions (Santos-Garcia & Jousselin 2027).

At the level of amplicon sequence variants (ASVs) our analysis identifies distinct, host-specific core microbiomes (Figure 2). The core microbiota of bugs feeding on the ancestral host plant, *C. corindum*, consists of a relatively even group of multiple taxa featuring Enterobacteriaceae and an unclassified Alphaproteobacteria ASV, whereas bugs on the introduced *K. paniculata* are dominated by a single unclassified Rhizobiaceae ASV. Remarkably, across these distinct host-associated cohorts, only a single ASV was conserved as a core member of both communities: a *Lactococcus* sequence variant. Lactic acid bacteria such as *Lactococcus* and *Lactobacillus* are frequent, functionally critical members of insect microbiomes (Vásquez et al 2012; Baffoni et al. 2026), often contributing to carbohydrate fermentation, altering gut pH, or defending against opportunistic pathogens. The consistent persistence of this specific *Lactococcus* ASV, regardless of the insect’s host plant or seasonal environment, suggests it may represent an obligate or beneficial symbiont for *J. haematoloma*.

Initial taxonomic classification revealed the presence of *Wolbachia*, a widespread intracellular endosymbiont known from several insect lineages (Werren 1997). We confirmed that *Wolbachia* is systemic across the head, gonads, and gut tissues of *J. haematoloma* using PCR on independently obtained insect tissue (Figure S2). This tissue distribution supports the presence of *Wolbachia* as an intracellular endosymbiont rather than as a resident of the gut lumen. However further investigation with microscopy will be necessary to definitively confirm that the bacterium is intracellular.

### Host plant is the primary determinant of community composition

Beta diversity comparisons delineate the gut microbiome into distinct communities driven by the host plant (Figure 4; Figure S4). The transition from the native balloon vine *C. corindum* to the introduced golden rain tree *K. paniculata* was likely rapid (Carroll & Boyd 1992), but these plants contain unique profiles of nutrients, saponins and cyanolipid toxins (Carroll et al. 1998). The segregation of microbial communities in association with these diets implies that the gut microbiota is shaped by the chemical environments created by feeding on these plants. These effects dominate other variables, including sex, geographic location or seasonality.

Alpha diversity analyses indicate that the complexity of the gut microbiome changes in response to both the host plant and the season, suggesting that metabolic requirements or microbial carrying capacities may differ. The host plant effect is most striking when examining the most abundant taxa (Figure 2). Bugs feeding on *C. corindum* maintain a more complex community composed largely of Enterobacteriaceae, with higher alpha diversity when compared to bugs feeding on *K. paniculata* (Figure 3A). Members of this family are common in hemipteran guts (Sandström et al. 2001; Li et al. 2022) and are implicated in metabolic roles, such as supplementing essential amino acids, synthesizing B-vitamins, or aiding in the enzymatic detoxification of plant defensive compounds (Cruden & Markovetz 1987; Douglas 1998). In contrast, the gut environment of bugs feeding on the introduced *K. paniculata* is dominated by a single unclassified Rhizobiaceae ASV, which dramatically reduces overall community evenness (Figure 3A). The dominance of this ASV suggests it may either be efficiently acquired from the plant’s environment or that it provides a localized metabolic advantage when processing *Koelreuteria* seeds or sap. While its precise metabolic activity remains unclear, many Rhizobiaceae are adapted to plant micro-environments and some species are capable of nitrogen fixation, sterol metabolism, or the degradation of complex polysaccharides (Bar-Shmuel et al. 2020), potentially helping the insect exploit this novel food source.

Beyond broad compositional shifts, consensus modeling of differentially abundant ASVs highlighted specific taxa associated with host and seasonal differences (Figure 5). In bugs feeding on the native host plant *C. corindum*, the gut microbiota was enriched with a *Caulobacter* ASV. Within the *K. paniculata* cohort, seasonal change was associated with the significant enrichment of two Bacilli species, *Pediococcus* and *Lactococcus*, during July as compared to January. The relative summer enrichment of this *Lactococcus* ASV, against the background of increased diversity in some lineages in July (Figure 3B, 4D), reinforces its potential functional importance as a consistent member of the soapberry bug microbiome core.

### Urbanization destabilizes community assembly

While environmental gradients like urbanicity often exert strong pressures on insect physiology, our results indicate that urbanization does not systematically alter taxonomic richness within soapberry bugs. Across the gradient of local green space, we did not observe obvious alpha diversity differences, paired with only subtle directional shifts in mean community composition (PERMANOVA). Instead, we found that local green space was associated with greater stability in the *J. haematoloma* gut microbiome. Tests for multivariate dispersion (PERMDISP) revealed a significant increase in variance among bugs collected from more urbanized sites, those with a lower proportion of local green space. This increased dispersion across both Bray-Curtis and UniFrac metrics indicates variance that can be caused by environmental stress (Zaneveld et al. 2017), or may be due to greater variance in microenvironmental factors in more urban sites. Rather than driving the microbial community toward a single, uniform “urban” state, human development appears to create more stochastic, variable microbiomes varying from individual to individual.

## Materials and Methods

### Sample collection and environmental metrics

Soapberry bugs (*Jadera haematoloma*) were collected from 15 sites in southern Florida during January and July 2019 (Table S1). Land cover data from the USGS National Land Cover Database (NLCD) for the surrounding 1-square kilometer area in 2019 was obtained from modelmywatershed.org (USGS 2019; Stroud Water Research Center 2017). The proportion of local green space was used as a metric of urbanicity (Nguyen & Rehan 2022; Borden et al. 2025) and was calculated as the total area (m^2^) of developed open space, deciduous forest, evergreen forest, mixed forest and cropland (NLCD codes 21, 41, 42, 43, 82) divided by 1 million m^2^ minus minus the area (m^2^) of open water (code 11). An urbanicity index was calculated for all sites using the first principle component from PCA on absolute area values for NLCD codes 11, 21-24, 31, 41-43, 52, 71, 81, 82, 90, and 95.

### Microbiota sequencing, ASV processing and taxonomic classification

Individual adult soapberry bugs were placed in 100% ethanol in the field and then stored at −20°C after confirmation of species determination in the lab with reference to Göllner-Scheiding (1979). The gut was dissected from stored individuals, and DNA was extracted using the Qiagen DNeasy Blood & Tissue Mini Kit. Extracted DNA was stored at −20°C. DNA samples were shipped on dry ice to the Centre for Comparative Genomics and Evolutionary Bioinformatics at Dalhousie University (Halifax, Canada). Samples were used to prepare paired-end 2 × 300-bp Illumina MiSeq libraries, with PCR amplification targeting variable regions 6-8 of the bacterial 16S ribosomal RNA gene.

Raw sequence quality was evaluated using FastQC v0.11.3 (Andrews 2010). Quality reports were aggregated using MultiQC v1.31 (Ewels et al. 2016) to generate comprehensive summary visualizations. All of the 180 samples demonstrated quality profiles with mean Phred scores >30 across the majority of positions. Data processing was conducted using the Qiime2 suite, v2021-04 (Bolyen et al. 2019). Illumina adapters and primer sequences were removed using the cutadapt plugin (Martin 2011). The --p-discard-untrimmed parameter was employed to exclude any read pairs lacking detectable primers. Amplicon sequence variants (ASVs) were inferred using DADA2 (Callahan et al. 2016), implemented as qiime dada2 denoise-paired. Denoising employed --p-trim-left-f 34 --p-trim-left-r 34 to remove low-quality bases and residual primer sequences from read termini, with --p-trunc-len-f 0 --p-trunc-len-r 0 (no length-based truncation) to maximize read length and overlap for paired-end merging. Post-denoising quality assessment via qiime feature-table summarize confirmed high sequence retention rates, with the majority of samples retaining >70% of input reads, indicating effective error correction without excessive data loss (Table S2). Sequence alignment, masking of variable regions and phylogenetic tree building was done via the qiime phylogeny align-to-tree-mafft-fasttree command to produce a mid-point rooted Newick-format phylogeny. Taxonomic assignments were performed using a pre-trained Naïve Bayes classifier based on the SILVA 138 99% reference database. Classification was performed using qiime feature-classifier classify-sklearn with 16 parallel threads, using a minimum posterior probability of 0.7.

### Screening for *Wolbachia*

To confirm the obligate nature of Wolbachia sequences identified in our 2019 taxonomic assignments, we performed PCR validation on a representative lab culture of *J. haematoloma* originally obtained from Lake Alfred, Florida in June 2024. The head, gut, and gonads (ovaries or testes) were dissected from 4 female and 4 male adults on individual sterile Petri dishes, and immediately moved to sterile microcentrifuge tubes for storage at −20°C. DNA was extracted using the Invitrogen PureLink Genomic DNA Mini Kit. The presence *Wolbachia* was inferred from amplification using the primer pairs W-Spec-16S-F/R (Werren & Windsor 2000) and ftsZ-F/R (Zha et al. 2014), which were designed specifically to target *Wolbachia*.

### Microbial diversity analysis

All statistical analyses were performed in R v4.6.1 (R Core Team 2026). Visualizations were generated using ggplot2 (Wickham 2016). Feature (ASV) counts were analyzed using the phyloseq package v1.56.0 (McMurdie & Holmes 2013) to manage count matrices and metadata, including taxonomic assignments. Data were filtered to remove counts from *Wolbachia*, mitochondria and chloroplasts, and a small number of human-associated Propionibacteriaceae. Samples with fewer than 1000 remaining reads were excluded from analysis, with retention of 144 samples. Raw counts were retained for differential abundance analysis, however sequencing depth was normalized for alpha diversity comparisons using the function phyloseq::rarefy_even_depth. Within-sample diversity was evaluated using counts of observed ASVs and Simpson’s index, as calculated by the phyloseq::estimate_richness function. To determine the significance of alpha diversity differences across categorical factors (such as host-plant and sampling month), we used the Wilcoxon rank-sum test. To examine the influence of continuous environmental factors, urbanicity and proportional green space, we constructed linear mixed-effects models using the lmer function from the lme4 package v2.0-6 (Bates et al. 2015). Collection site was included as a random intercept in the mixed models to account for local non-independence.

The phyloseq::distance function was used to calculate sample distances according to the Bray-Curtis, unweighted UniFrac, and weighted UniFrac metrics. Global community variance was visualized using principal coordinates analysis (PCoA). To account for unbalanced field sampling, some hypothesis testing was restricted to data subsets. The effect of the host plant was evaluated within July samples, while the effect of seasonality (month) was evaluated within the cohort of bugs collected from *K. paniculata*. Permutational multivariate analysis of variance (PERMANOVA) was conducted on these subsets using the adonis2 function in the vegan package v2.7-6 (Oksanen et al. 2026) with 1000 permutations and marginal sums of squares (by = “margin”). To determine whether urbanization primarily drove directional community shifts or altered community heterogeneity, we assessed multivariate homogeneity of group dispersions (PERMDISP) using the vegan::betadisper and permutest functions with 10^5^ iterations, categorizing the continuous green space metric into statistical tertiles (low, medium, high green space).

To visualize global community variance across host plants and seasons, we performed generalized linear model principal component analysis (GLM-PCA) using the glmpca package v0.2.0 (Townes & Street 2020). We specified a negative binomial error distribution to account for overdispersion and to normalize differing sequencing depths as a latent covariate.

To identify the differential abundance of taxa associated with metadata factors, we utilized three methods, including ANCOM-BC2 (Lin & Peddada 2024), ALDEx2 v1.44.0 (Fernandes et al. 2014), and MaAsLin3 v2.0.2 (Teichman & Willis 2026). For two-level categorical factors ADLEx2 significance was determined using Welch’s t-test with Benjamini-Hochberg false discovery rate correction (FDR < 0.05) and an absolute expected effect size threshold of > 0.5.

To identify the differential abundance of taxa associated with metadata factors, we utilized a multi-algorithm consensus approach combining ANCOM-BC2 (Lin & Peddada 2024), ALDEx2 v1.44.0 (Fernandes et al. 2014), and MaAsLin3 v1.5.7 (Nickols et al. 2026) applied to the same data subsets described above. For two-level categorical factors, ALDEx2 significance was determined using Welch’s t-test. For continuous environmental factors (proportional green space), values were converted to standardized Z-scores and evaluated via generalized linear modeling (`aldex.glm`). Across all algorithms, significance was defined using Benjamini-Hochberg false discovery rate correction. Taxa were considered robust differential drivers if they were identified as statistically significant by at least two of the three algorithms.

## Data availability

Code and data involved in this analysis are available online at https://github.com/aphanotus/Jhae.microbiome

## Acknowledgements

Research reported in this publication was supported by the Colby College Division of Natural Sciences and by an Institutional Development Award (IDeA) from the National Institute of General Medical Sciences of the National Institutes of Health under grant number P20GM0103423.

