## Supplemental Materials for "Host specificity, seasonality, and urbanicity shape the gut microbiome of a synanthropic insect"

for

Noyes, et al.

Supplemental Tables appear in the accompanying spreadsheet file.

#### Supplemental Tables

**Table S1: Collection sites.**

| Municipality | Latitude | Longitude | Host plant | Sample size in January | Sample size in July | Proportion of green space | Urbanicity index |
|---|---|---|---|---|---|---|---|
| Lake Wales FL | 27.9594 | -81.5941 | *K. paniculata* | 2 | 3 | 0.7579 | -3.385 |
| Lake Wales FL | 27.8438 | -81.5853 | *K. paniculata* | 0 | 25 | 0.4180 | -2.621 |
| Islamorada FL | 24.9501 | -80.5957 | *C. corindum* | 0 | 5 | 0.0539 | -2.517 |
| Bartow FL | 27.9549 | -81.7708 | *K. paniculata* | 5 | 1 | 0.2041 | -1.988 |
| Lakeland FL | 28.0927 | -81.9536 | *K. paniculata* | 0 | 4 | 0.1697 | -0.014 |
| Lake Alfred FL | 28.0918 | -81.7231 | *K. paniculata* | 6 | 14 | 0.4207 | 1.107 |
| Winter Haven FL | 28.0433 | -81.7615 | *K. paniculata* | 3 | 0 | 0.2931 | 1.312 |
| Lake Bentley FL | 28.0211 | -81.9322 | *K. paniculata* | 3 | 0 | 0.5783 | 1.403 |
| Fort Lauderdale FL | 26.1884 | -80.1720 | *K. paniculata* | 10 | 9 | 0.0806 | 1.697 |
| Lakeland FL | 28.0334 | -81.9552 | *K. paniculata* | 6 | 10 | 0.2920 | 1.758 |
| Lehigh Acres FL | 26.6173 | -81.6897 | *K. paniculata* | 5 | 0 | 0.4459 | 1.763 |
| Delray Beach FL | 26.4749 | -80.0912 | *K. paniculata* | 6 | 19 | 0.0929 | 1.916 |
| Kendall Lakes FL | 25.7139 | -80.4203 | *K. paniculata* | 8 | 0 | 0.2868 | 2.102 |

**Table S2: DADA2 Denoise samples statistics.** The number of input reads, those passing quality filter and their percentage, the number denoised and merged and their percentage, the number of non-chimeric sequences and their percentage.

This table is very large. Please refer to the accompanying spreadsheet.

**Table S3: Core microbiome in each host plant cohort.**

| Host plant | ASV ID | Mean abundance | Prevalence | Phylum | Class | Order | Family | Genus |
|---|---|---|---|---|---|---|---|---|
| *C. corindum* | 02502141f0a1bf07cc6d6c60b94433f5 | 0.278790 | 1 | Proteobacteria | Gammaproteobacteria | Enterobacterales | Enterobacteriaceae | NA |
| *C. corindum* | 5f9403f34de2d6658df09b913aa13184 | 0.242246 | 1 | Proteobacteria | Alphaproteobacteria | NA | NA | NA |
| *C. corindum* | 7364a9d6bfb4a993fabfa2e22252a18d | 0.230191 | 1 | Proteobacteria | Gammaproteobacteria | Enterobacterales | Enterobacteriaceae | NA |
| *C. corindum* | 8084aa68823c09679d87c31776e22478 | 0.150427 | 1 | Firmicutes | Bacilli | Lactobacillales | Streptococcaceae | Lactococcus |
| *K. paniculata* | 9f8c37a25eca1c8caec4c05d14cd7cdf | 0.661537 | 0.992 | Proteobacteria | Alphaproteobacteria | Rhizobiales | Rhizobiaceae | NA |
| *K. paniculata* | 8084aa68823c09679d87c31776e22478 | 0.135416 | 0.841 | Firmicutes | Bacilli | Lactobacillales | Streptococcaceae | Lactococcus |

**Table S4: Sample alpha diversity metrics.**

This table is very large. Please refer to the accompanying spreadsheet.

**Table S5: Alpha test statistics.** Binary categorical factors were tested using the Wilcoxon rank sum test, producing a W statistic. Continuous factors were modeled in linear mixed effects models with collection site as a random intercept, producing t statistics. Associated p-values are listed without correction.

| metric | factor | W | p | |
|---|---|---|---|---|
| observed_ASVs | sex | 2149.5 | 0.07717 | |
| shannon | sex | 2500.5 | 0.7176 | |
| simpson | sex | 2593.5 | 0.9952 | |
| observed_ASVs | morph | 2074.0 | 0.4926 | |
| shannon | morph | 2112.0 | 0.6029 | |
| simpson | morph | 2108.0 | 0.5909 | |
| observed_ASVs | host_plant | 316.0 | 0.06921 | |
| shannon | host_plant | 352.0 | 0.01435 | * |
| simpson | host_plant | 356.0 | 0.01177 | * |
| observed_ASVs | month | 1491.0 | 0.0005031 | *** |
| shannon | month | 1306.5 | 0.00001957 | *** |
| simpson | month | 1337.5 | 0.00003538 | *** |
| **metric** | **factor** | **t** | **p** | |
| observed_ASVs | urbanicity | -0.846 | 0.423 | |
| shannon | urbanicity | -1.544 | 0.182308 | |
| simpson | urbanicity | -1.553 | 0.174039 | |
| observed_ASVs | green_space | 0.534 | 0.60365 | |
| shannon | green_space | 0.534 | 0.60365 | |
| simpson | green_space | -1.369 | 0.198404 | |

#### Table S6: PERMANOVA test statistics.

| Distance matrix | factors | df | SS | $R^2$ | F | p | |
|---|---|---|---|---|---|---|---|
| Bray-Curtis | sex | 1 | 0.1469 | 0.00491 | 0.7919 | 0.5005 | |
| | morph | 2 | 0.2861 | 0.00956 | 0.7714 | 0.554446 | |
| | host_plant | 1 | 2.0693 | 0.06914 | 11.1585 | 0.000999 | *** |
| | month | 1 | 0.3815 | 0.01275 | 2.0574 | 0.090909 | . |
| | green_space | 1 | 0.8789 | 0.02937 | 4.7394 | 0.001998 | ** |
| | Residual | 137 | 25.4065 | 0.84892 | | | |
| | Total | 143 | 29.928 | 1 | | | |
| unweighted UniFrac | sex | 1 | 0.412 | 0.00937 | 1.4688 | 0.080919 | . |
| | morph | 2 | 0.741 | 0.01687 | 1.3215 | 0.105894 | |
| | host_plant | 1 | 1.819 | 0.04138 | 6.4842 | 0.000999 | *** |
| | month | 1 | 1.542 | 0.03509 | 5.4979 | 0.000999 | *** |
| | green_space | 1 | 0.994 | 0.0226 | 3.5418 | 0.000999 | *** |
| | Residual | 137 | 38.434 | 0.87437 | | | |
| | Total | 143 | 43.956 | 1 | | | |
| weighted UniFrac | sex | 1 | 0.00017 | 0.00009 | 0.0137 | 0.94805 | |
| | morph | 2 | 0.01561 | 0.0087 | 0.6337 | 0.56543 | |
| | host_plant | 1 | 0.02669 | 0.01488 | 2.1675 | 0.1009 | |
| | month | 1 | 0.00852 | 0.00475 | 0.6918 | 0.46853 | |
| | green_space | 1 | 0.04483 | 0.025 | 3.6411 | 0.04296 | * |
| | Residual | 137 | 1.6868 | 0.94068 | | | |
| | Total | 143 | 1.79317 | 1 | | | |
| Bray-Curtis | host_plant | 1 | 2.6586 | 0.14451 | 14.865 | 0.000999 | *** |
| | Residual | 88 | 15.7391 | 0.85549 | | | |
| | Total | 89 | 18.3977 | 1 | | | |
| unweighted UniFrac | host_plant | 1 | 1.7687 | 0.07362 | 6.9929 | 0.000999 | *** |
| | Residual | 88 | 22.2572 | 0.92638 | | | |
| | Total | 89 | 24.0258 | 1 | | | |
| weighted UniFrac | host_plant | 1 | 0.03196 | 0.03447 | 3.1414 | 0.04595 | * |
| | Residual | 88 | 0.89516 | 0.96553 | | | |
| | Total | 89 | 0.92712 | 1 | | | |
| Bray-Curtis | month | 1 | 0.4096 | 0.01536 | 2.1375 | 0.1039 | |
| | Residual | 137 | 26.2553 | 0.98464 | | | |
| | Total | 138 | 26.6649 | 1 | | | |
| unweighted UniFrac | month | 1 | 1.679 | 0.04059 | 5.7962 | 0.000999 | *** |
| | Residual | 137 | 39.683 | 0.95941 | | | |
| | Total | 138 | 41.361 | 1 | | | |
| weighted UniFrac | month | 1 | 0.0096 | 0.00548 | 0.7547 | 0.4236 | |
| | Residual | 137 | 1.742 | 0.99452 | | | |
| | Total | 138 | 1.7516 | 1 | | | |

**Table S7: ANCOM-BC2 Results.**

This table is very large. Please refer to the accompanying spreadsheet.

**Table S8: ALDEx2 Results.**

This table is very large. Please refer to the accompanying spreadsheet.

**Table S9: MaAsLin3 Results.**

This table is very large. Please refer to the accompanying spreadsheet.

**Table S10: Consensus of differential abundance analysis methods.**

This table is very large. Please refer to the accompanying spreadsheet.

**Table S11: PERMDISP Results.**

| Distance matrix | factors | Df | SS | MSS | F | Pr(>F) | |
|---|---|---|---|---|---|---|---|
| Bray-Curtis | Groups | 2 | 1.4526 | 0.72629 | 7.3334 | 0.00104 | ** |
| | Residuals | 136 | 13.4691 | 0.09904 | | | |
| unweighted UniFrac | Groups | 2 | 0.4246 | 0.212289 | 7.086 | 0.00155 | ** |
| | Residuals | 136 | 4.0744 | 0.029959 | | | |
| weighted UniFrac | Groups | 2 | 0.10743 | 0.053715 | 6.3726 | 0.00223 | ** |
| | Residuals | 136 | 1.14635 | 0.008429 | | | |

### Supplemental Figures

#### Figure S1

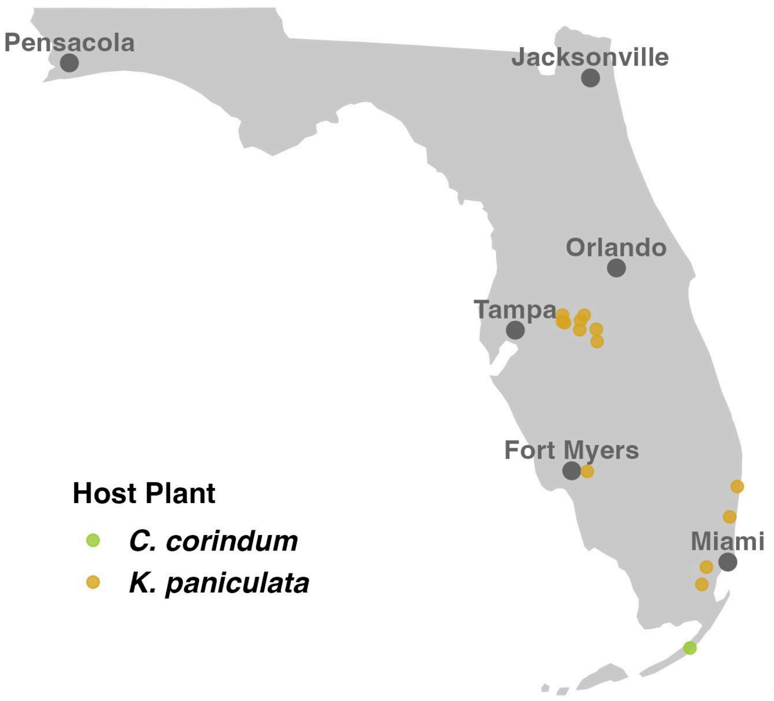

**Figure S1: Map of collections sites in this study.**

**Figure S2**

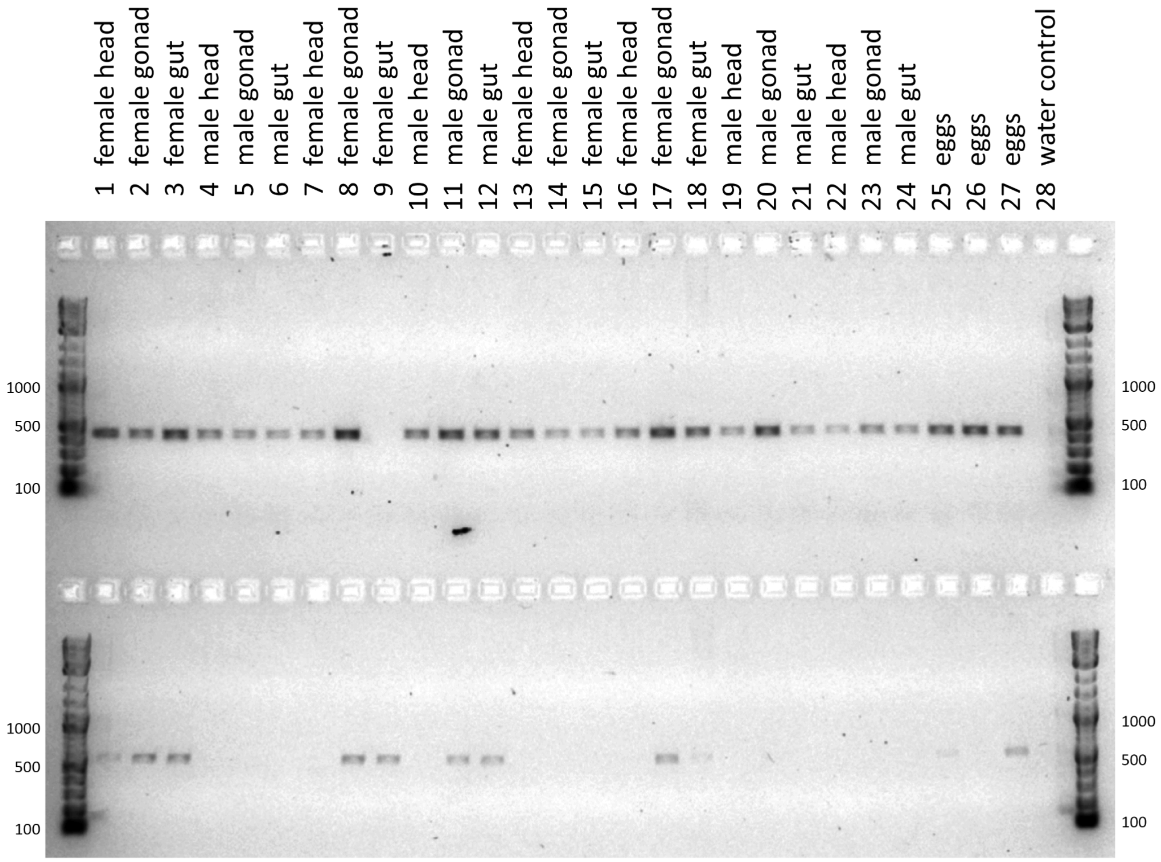

**Figure S2: PCR screen for the presence of *Wolbachia* in *J. haematoloma* tissue.** Template DNA from the head, gonads and gut of 4 female and 4 male adults, as well as 3 separate pools of eggs were used in PCRs targeting *Wolbachia*-specific sequences. Each template was tested with two primer pairs. PCR rxn conditions. Primers W-Spec-16S-F/R (Werren & Windsor 2000; top row) produce an expected product of 438 bp. All samples except one female gut sample produced the expected product. Primers ftsZ-F/R (Zha et al. 2014; bottom row) produce an expected product of 570 bp. Only some individuals appeared to be positive for *Wolbachia* based on this primer pair. All sampled tissues showed evidence of *Wolbachia* in at least one individual. Negative (water) controls were consistently negative for amplification with both primers pairs. Product sizes were determined by comparison to be New England BioLabs 1-kb Plus DNA ladder (outside lanes).

### Figure S3

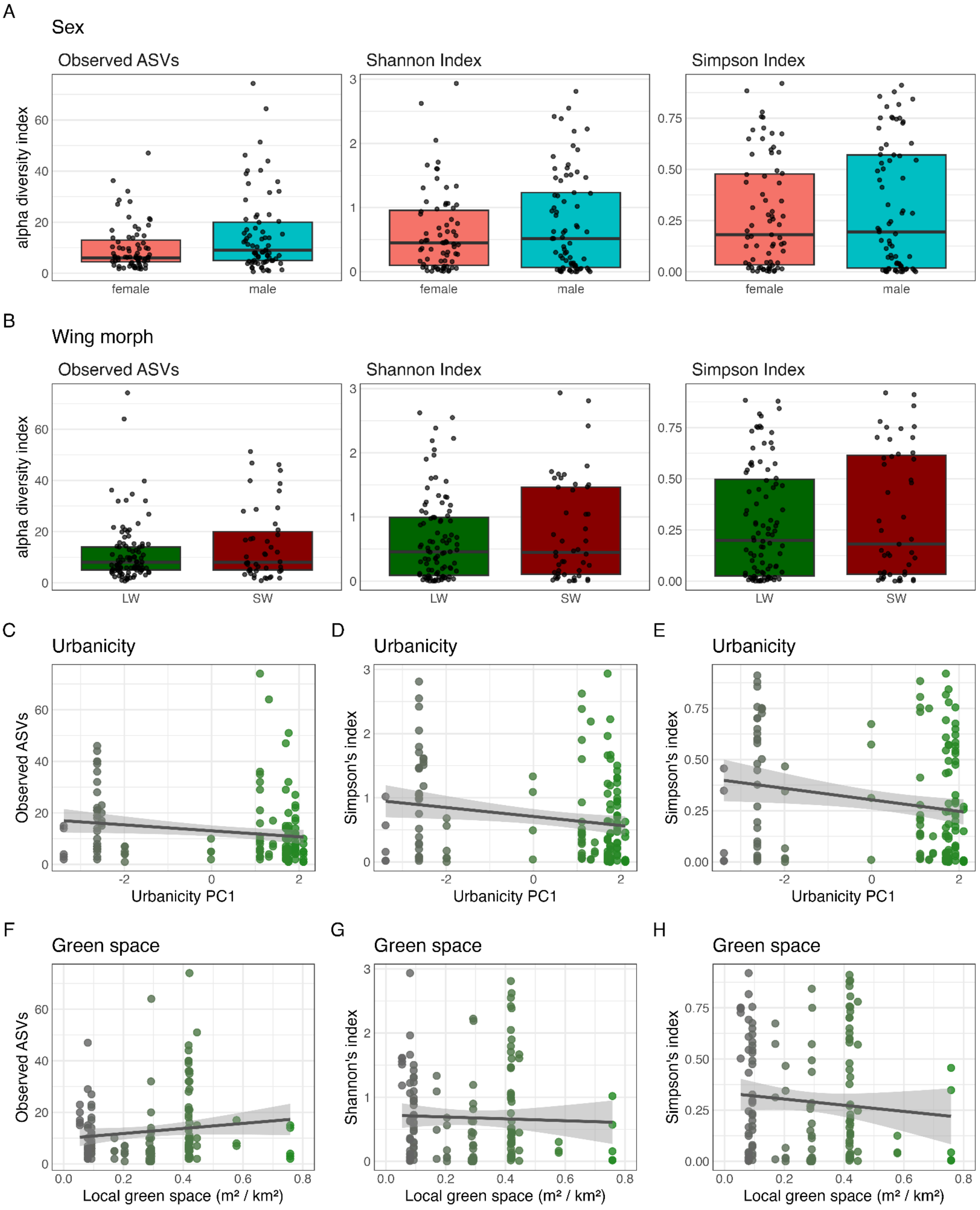

**Figure S3: Non-significant factors in *J. haematoloma* gut microbiota alpha diversity. (A-B)** Box plots displaying within-sample taxonomic richness and diversity. Each panel plots observed ASVs, Shannon index and Simpson's index. Center lines represent the median, box limits indicate the upper and lower quartiles, and whiskers extend to 1.5 times the interquartile range. Statistical significance was tested using Wilcoxon rank-sum tests (Table S5). Factors with significant association to alpha diversity are shown in Figure 3. **(A)** Alpha diversity comparisons by sex. Females: n = 71; males = 73. **(B)** Alpha diversity comparisons between wing morphs: long-winged (LW, n = 95) and short-winged (SW, n = 47) adults. **(C-H)** Continuous factors are shown in scatterplots with a best-fit line (black) bounded by a 95% confidence interval (gray fill, n = 139). **(C-E)** Urbanicity, calculated as the first principle component from an analysis of land cover data at each collection site. **(F-H)** Proportion of local green space.

**Figure S4**

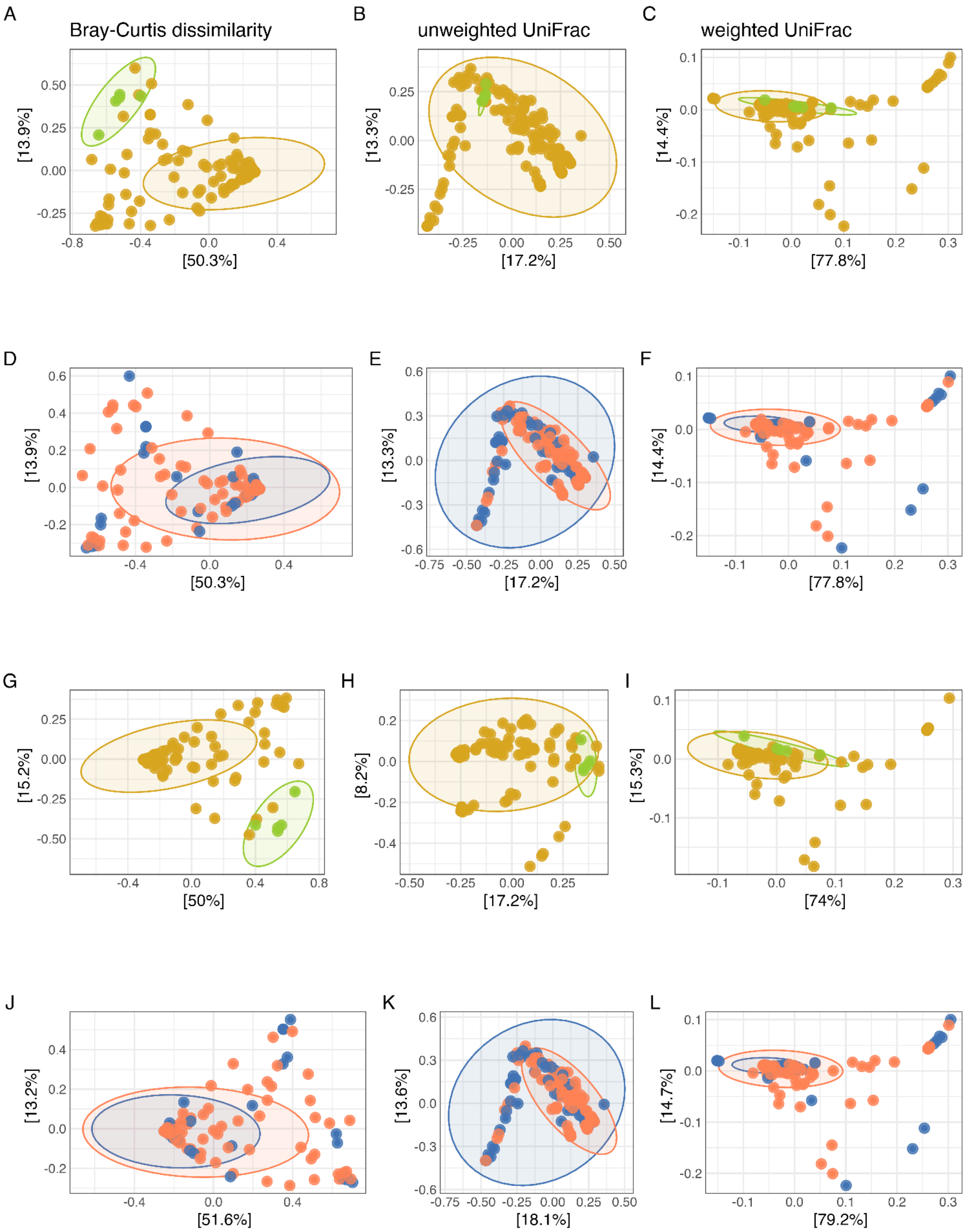

**Figure S4: Principle coordinate analysis (PCoA) plots.** Panels display the first and second PCoA axes for Bray-Curtis (A, D, G, J), unweighted UniFrac (B, E, H, K), and weighted

UniFrac (C, F, I, L) distances. Host plant association is highlighted in panels A-C and G-I with samples from the ancestral host plant *Cardiospermum corindum* in green and the introduced host plant *Koelreuteria paniculata* in orange. Samples collected in different seasons are color-coded in panels D-F and J-L, with January samples in dark blue and July samples in light red. PCoA was initially applied to the complete dataset (A-F). To address the unbalanced nature of sample collection, we also examined the effects of these factors in data subsets. Host plant effects were examined only among July samples (G-I) and seasonal effects were examined only among samples collected from *K. paniculata* (J-L). Shaded ellipses represent 95% confidence intervals assuming a multivariate distribution (`stat_ellipse`).
